# A system biology approach reveals cellular pathway differences between follicular thyroid carcinoma and follicular thyroid adenoma

**DOI:** 10.1101/480632

**Authors:** Md. Ali Hossain, Tania Akter Asa, Md. Mijanur Rahman, Julian M.W. Quinn, Fazlul Huq, Mohammad Ali Moni

**Author notes:** Corresponding Author Email address (Md. Ali Hossain), (Mohammad Ali Moni).

## Abstract

Pathogenic mechanisms that underlie malignant follicular thyroid carcinoma (FTC) development are poorly understood. To identify key genes and pathways driving malignant behaviour we employed a system biology-based integrative analyses comparing FTC transcriptomes with a similar but benign lesion, follicular thyroid adenoma (FTA). We identified differentially expressed genes (DEGs) in microarray gene expression datasets (n=52) of FTCs and FTA tissues. Pathway analyses of DEGs using gene ontology (GO) and Kyoto Encyclopedia of Genes and Genomes (KEGG) resources revealed significant pathways, and pathway hub genes using protein-protein interactions (PPI). We identified 598 DEGs (relative to FTAs) in FTCs and 12 significant pathways with altered expression in FTC. 10 GO groups were significantly connected with FTC-high expression DEGs and 80 with low-FTC expression. PPI analysis identified 12 potential hub genes based on degree and betweenness centrality. Moreover, 10 transcription factors (TFs) were identified that may underlie DEG expression as well as a number of microRNA (miRNAs). Thus, we identified DEGs, pathways, TFs and miRNAs that reflect molecular mechanisms differing between FTC and benign FTA. These may constitute biomarkers that distinguish these lesions and, given the similarities and common origin of the lesions, they may also be indicators of malignant progression potential.

## 1. Introduction

Thyroid cancers are the most common type of endocrine malignancy, although they have a relatively low mortality rate compared to most other common metastatic diseases. The United States had 56,460 new diagnoses of thyroid cancer and 1,780 related deaths reported in 2012 [18]. Its incidence is also rising globally at about 5% per year, although some of this increase may be due to improved detection, and it notably affects those in the 20 to 34 year age range [1]. Thyroid cancers include several major types including papillary thyroid carcinomas, medullary thyroid carcinoma, anaplastic thyroid carcinoma and follicular thyroid carcinomas (FTCs) [3]; FTC is one of the more aggressive types, although it accounts for a minority (14%) of total thyroid cancers [34].

The causes and cellular processes that give rise to FTC and control these tumors behaviour are poorly understood; because of this, these cancers have few effective treatment options [37]. There is, therefore, a great need to understand the mechanisms that drive development and progression in FTC to identify new approaches to detection, estimate risk of progression and find new therapies. In addition, differential diagnosis of FTC is problematic as it can be difficult to distinguish from follicular thyroid adenoma (FTA), a benign and non-invasive lesion. For this reason, molecular markers that distinguish FTC and FTA (and other types of thyroid lesions) are much sought. Thus, Wojtas *et al.* conducted a gene expression comparison of FTC and FTA lesions which identified potential markers that can distinguish FTC from FTA with a sensitivity and specificity of 78% and 80%, respectively. [51]. We can also use this dataset to identify pathways with different levels of activity in benign and aggressive thyroid tumours (here, FTA and FTC) that may reflect important molecular mechanisms that underlie their behaviour.

For such pathway studies our starting point is the differential expression of genes (DEGs) between these lesions. We can then use analytic tools to study these DEGs and discover pathways, and pathway hub genes that may affect cell functions and thereby underlie their tumour morbidity, growth and invasion [28, 14, 30]. The repertoire of candidate pathway factors can be extended using approaches such as gene ontology analysis and protein-protein interactions (PPI) studies.

To identify such DEGs, microarray gene expression profiling is widely used [11] and several such studies have been performed for thyroid cancer subtypes [40]. For example, Huang et al. [16] studied gene expression profiles of thyroid cancers, although not functional interactions between gene products. To understand better the underlying molecular mechanisms and identify critical biomolecules, integrative analysis within a gene network context is needed [38, 29, 39]. These may lead to candidate FTC biomarkers (the focus of Wojtas *et al.*) but our main interest is to obtain key or hub genes distinguishing malignant FTC from benign FTA as these may be potential therapeutic targets. Such a systems biology approach integrating network statistical and topological analyses of experimental datasets can thus clarify disease mechanisms [22]. Thus, we used such a systems approach (Figure 1) to identify FTC molecular signatures at the miRNA, mRNA, and protein levels that differ from that of FTAs. Such a comparison should be more meaningful than a comparison of FTC to normal thyroid tissue as the tumours are similar, yet differ markedly in their potential for invasion metastasis.

**Figure 1:**
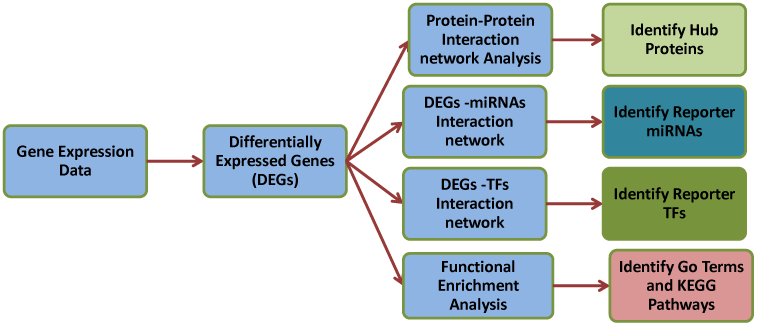
The multi-stage analysis methodologies employed in this study. Gene expression datasets related to FTC and FTA tissue were collected from the NCBI Gene Expression Omnibus (NCBI-GEO) database and statistically analyzed using GEO2R to identify DEGs. Four types of functional enrichment analyses of DEGs were then performed to identify significantly enriched pathways. Thus, we constructed protein-protein interaction networks around DEGs topological analyses to identify putative pathways hub proteins, identified possible micro-RNA (miRNA) and transcription factor (TF) interactors, and used Gene Ontology annotation terms to provide pathways enrichment. TF and miRNA studies employed JASPAR and miRTarbase databases, respectively. DEGs were integrated with those networks and higher degree and betweenness centrality were used to designate TFs and miRNAs as the reporter transcriptional regulatory elements. The target DEGs of reporter miRNAs and TFs were subjected to pathway enrichment analyses.

## 2. MATERIALS AND METHODS

In this study, the multi-step analysis method we developed and applied is shown in Fig. 1. We statistically analyzed gene expression datasets to identify the DEGs and their regulatory patterns. We employed these DEGs to identify enriched pathways, biological processes and annotation terms (i.e., Gene Ontology terms) by using functional enrichment methods. Then, to identify reporter biomolecules, we integrated the intermediate analysis results with biomolecular networks.

### Dataset Employed and Statistical Methods Used

We obtained the gene expression data of FTC and FTA (GSE82208) for our study from the NCBI Gene Expression Omnibus (GEO) (http://www.ncbi.nlm.nih.gov/geo/) [4, 51]. This dataset contained analyses of RNA from frozen tumour tissue specimens from 27 FTC and 25 FTA lesions using Affymetrix human genome U133 (Plus 2.0) arrays. The lesions were diagnosed histologically, many with a second diagnosis to confirm, when paraffin embedded material was available [51]. Gene expression analysis using microarrays is a widely used method to develop and refine the molecular determinants of human disorders. We used data using these technologies to analyze the gene expression profiles of FTC and FTA. To identify the DEGs between FTC samples and FTA, t-test method was used by using the Limma package [47]. To identify the up-regulated genes, we used the condition of *p − value* < .05 and *logFC* > −2 (FC, fold change) and for identifying the down-regulated genes, *p − value* < .05 and *logFC* < − 2 were used. All identified up-regulation genes and down-regulation genes were considered as DEGs. We applied the topological and neighborhood based benchmark methods to find gene-gene associations. A gene-gene network was constructed by using the gene-gene associations, where the nodes in the network represent gene [53, 29]. This network can also be characterized as a bipartite graph. These topological and neighborhood based benchmark methods were adopted from our previous studies [30].

The common neighbours are the based on the Jaccard Coefficient method, where the edge prediction score for the node pair is as [31]:

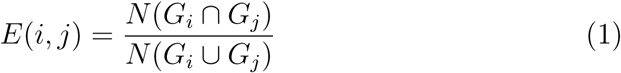

where *G* is the set of nodes and *E* is the set of all edges. We used R software packages “comoR” [28] and “POGO” [30] to cross check the genes-diseases associations.

### Functional Enrichment of Gene Sets

We performed gene ontology and pathway analysis on identified up-regulation genes and down-regulation genes using DAVID bioinformatics resources (https://david-d.ncifcrf.gov/) (version v6.8) [46] to get further insight into the molecular pathways that differ between FTC and FTA. In these analyses, GO and KEGG pathway databases were used as annotation sources. Enrichment results showing an adjusted *p − value* < 0.05 were considered significant.

### Construction and Analysis of Protein-Protein Interaction (PPI) Sub-networks

The PPI network was first constructed with the DEGs and analyzed using STRING [48] a web-based visualization software resource. The constructed PPI network was represented as an undirected graph, where nodes represent the proteins and the edges represent the interactions between the proteins. To construct the PPI network from the STRING database (http://string-db.org) [48], we used database data, data mined from PubMed abstract text, Co-expression, gene fusion and Neighborhood as active interaction sources and a combined score that is greater than 0.4 was set as the level of significance. The PPI network was then visualized and analyzed using Cytoscape (v3.5.1) [45, 32]. Then, topological analysis was applied to identify highly connected proteins (i.e., hub proteins) through the Cyto-Hubba plugin [7] where betweenness centrality and higher degree were employed. After then, the top three modules (i.e., the three most highly interconnected protein clusters) in the PPI subnetwork were identified using the MCODE plug-in [7]. Finally, these modules were further analyzed and characterized using enrichment analyses by NetworkAnalyst [52, 13]. The KEGG pathway enrichment analysis of the PPI networks involved DEGs were performed by NetworkAnalyst [52, 13].

### Identifying TFs and miRNAs that Influence Expression of Candidate Genes

To identify TFs and miRNAs that affect transcript levels around which significant changes occur at the transcriptional level, we obtained experimentally verified TF-target genes from the JASPAR database [20] and miRNA-target gene interactions from TarBase [43] and miRTarBase [15] by using NetworkAnalyst tools [52] where betweenness centrality and higher degree filters were used. Currently, there were many techniques to measure the topological properties. We used the Degree Centrality (DC) and Betweenness Centrality (BC) to find out networks topological properties. We can define the DC of a node v in a network as the total number of nodes which are directly connected to node v in that network. The definition can also be written as follows:

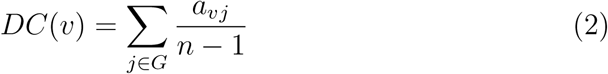

Whereas, n represents total number of nodes in the network and avj represents that node v and node j are directly connected. In the case of Betweenness Centrality (BC), the total number of times of node v appearing in the shortest path between other nodes are quantified. It is also defined as follows:

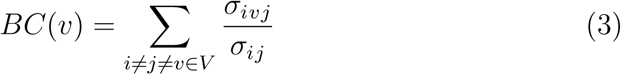

Where *σ*_*i*_*j*__ = total number of shortest paths from node i to node j, and *σ*_*ivj*_ = total number of paths through node v.

## 3. Results

### 3.1. Results of DEG analyses

#### Transcriptomic signatures: Differentially expressed genes

FTA and FTC tissue gene expression patterns were analysed using oligonucleotide microarrays from the NCBI GEO (http://www.ncbi.nlm.nih.gov/geo/query/acc.cgi?acc=G [51]. Gene expression profiling was performed in 27 malignant FTCs and 25 FTAs. 598 genes were differentially expressed (*p* < 0.05, > 1.0 log2 fold change) relative to FTAs, of which 465 genes were significantly lower expression and 133 genes were higher expression levels in FTC lesions (see Additional file 1:Table S1).

### 3.2. Pathway and functional correlation analysis

Combining large scale, state of the art transcriptome and proteome analysis, we performed a regulatory analysis to gain further insight into the molecular pathways associated with the FTC and predicted links to pathways that differ relative to the benign FTAs. DEGs and pathways were analysed using KEGG pathway database (http://www.genome.jp/kegg/pathway.html) and functional annotation tool DAVID v 6.8 (http://niaid.abcc.ncifcrf.gov) to identify overrepresented pathway groups and it was observed that 4 pathways were associated with DEGs with higher expression in FTC compared to FTA, namely the “One carbon pool by folate” pathway, p53 signaling pathway, Cell cycle and Progesterone-mediated oocyte maturation signaling pathway. Fold enrichment, adjusted p-values and genes associated with these pathways are presented in Table 1(a) A.

**Table 1:**
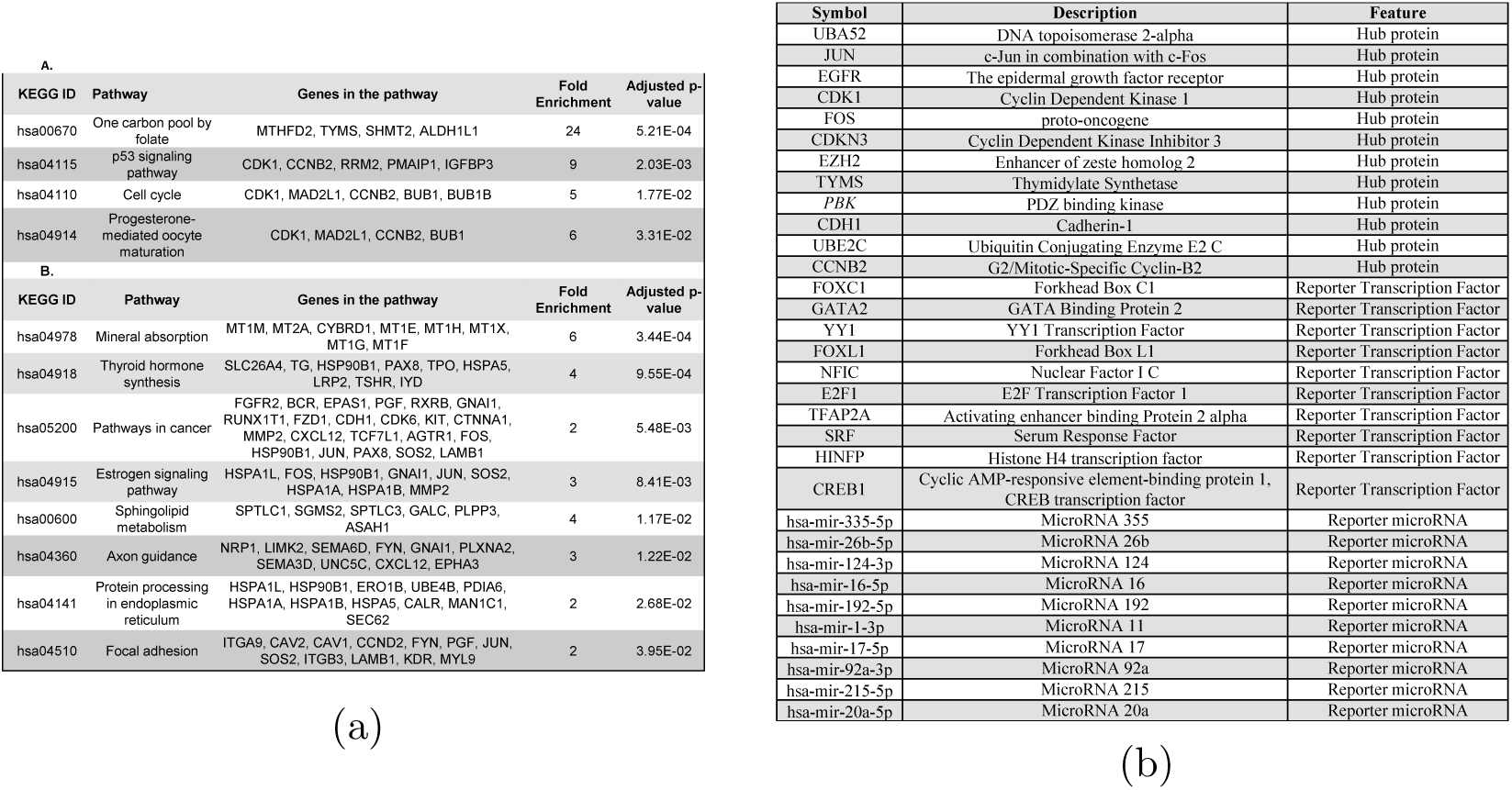
(a) Part A shows pathways associated with the genes showing higher expression in FTC than FTA. This set showed an enrichment of four KEGG pathways, annotated using DAVID. Part B shows pathways associated with genes showing lower expression levels in FTC. This set showed in an enrichment of eight KEGG pathways using DAVID. Fold enrichment, adjusted p-values and genes associated with these pathways are indicated. (b) Summary of reporter biomolecules (including putative hub genes, TFs, and miRNAs) in that differ between FTC and FTA.

It was also observed that 8 significantly over-represented pathways including Mineral absorption, Thyroid hormone synthesis, Pathways in cancer, Estrogen signaling, Sphingolipid metabolism, Protein processing in endoplasmic reticulum, Axon guidance and Focal adhesion pathways (Table 1(a) B) were identified. These are associated with the DEGs expressed at lower levels in FTC lesions compared to FTA. We then performed GO analysis using DAVID to obtain further insight into the molecular roles and biological function of DEGs identified in this study. From this analysis, 9 GO groups were associated with DEGs that were more highly expressed in FTC (see Table 2(a)). Reflecting their greater number, the genes with lower expression in FTC compared to FTA were associated with 80 GO groups; the 10 most significant GO groups are presented in Table 2(b). Fold enrichment, adjusted p-values and genes associated with these ontologies are also indicated.

**Table 2:**
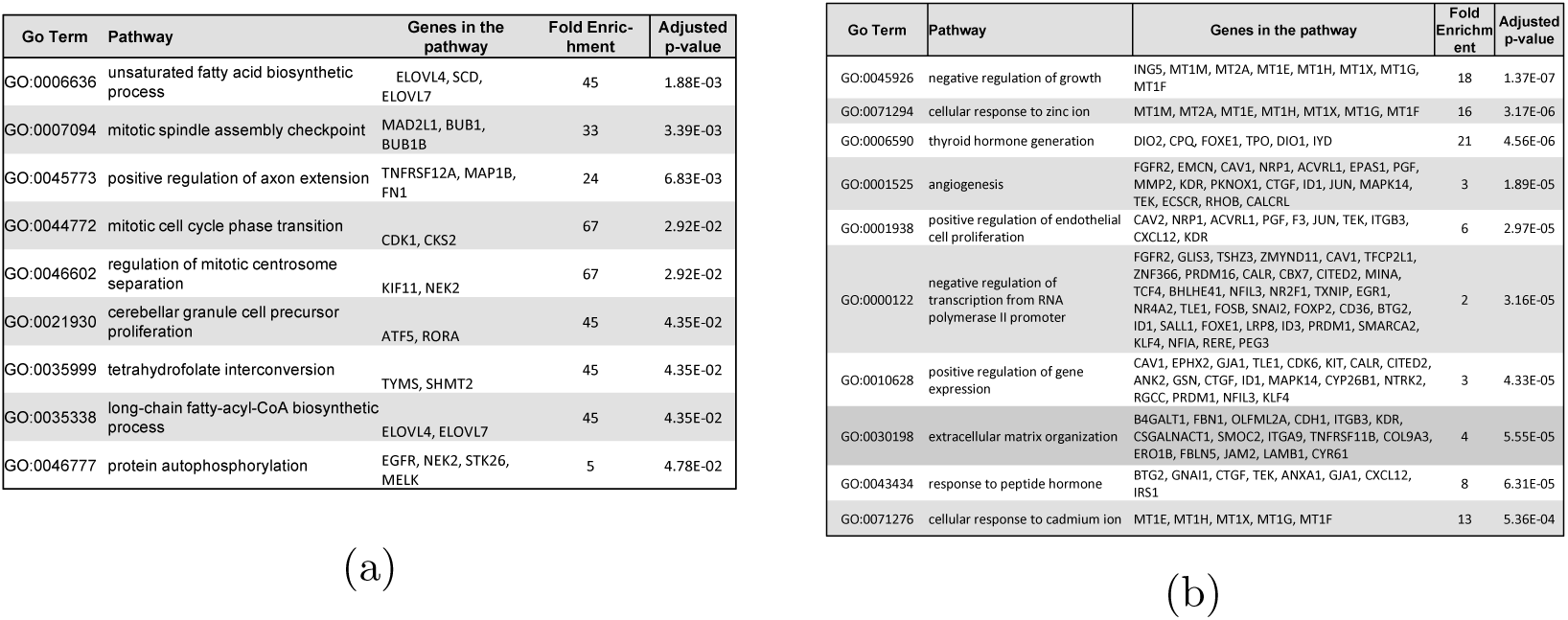
(a) Gene ontologies associated with genes with higher expression in FTC. (b) Gene ontologies associated with genes with lower expression in FTC compared to FTA. The 10 most significant biological functional ontologies are indicated and are those associated with these significant genes. Fold enrichment, adjusted p-values and genes associated with these ontologies are presented in these tables.

### 3.3. Proteomic signatures: Hub target proteins from PPI analysis

A PPI network was constructed using the DEGs identified in this study (Fig. 2A) using the STRING package. Topological analyses using STRING as well as further analysis by Cytoscape’s Cyto-Hubba plugin identified the most significant hub proteins, which were identified as gene products of TOP2A, JUN, EGFR, CDK1, FOS, CDKN3, EZH2, TYMS, PBK, CDH1, UBE2C and CCNB2. The simplified PPI network for most significant hub genes were constructed by using Cyto-Hubba plugin and are shown in fig. 2B.

**Figure 2:**
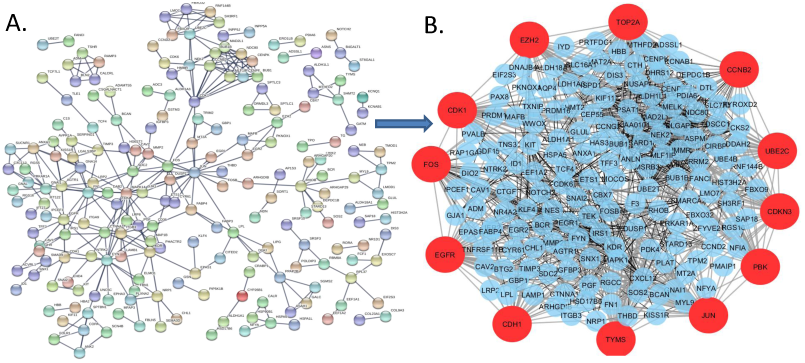
A. The protein-protein interaction network constructed using the DEGs identified in the FTC/FTA comparison. B. The protein-protein interaction(PPI) network identified in FTC/FTA dataset DEGs showing the most significant hub genes on the periphery (red)

### 3.4. Enrichment Analysis of the Modules found in PPI of DEG

The PPI network was analyzed by using MCODE plug-in in Cytoscape (version 3.5.1), and top three modules were selected (Fig. 3). The enrichment analysis of the top three modules was analyzed by using DAVID. Module 1 represented a set of biological process significantly enriched for ATP binding, Cell division, Cell proliferation; associated enriched cellular components were Kinetochore, Condensed chromosome kinetochore (both of which relate to cell division) and Nucleoplasm and Cytoplasm which are very general categories. Molecular functions significantly enriched in module 1 included chromatin binding and protein kinase activity (Table 3 (a)). Module 2 showed significantly enriched biological process positive regulation of Protein phosphorylation and Protein autophosphorylation; these significantly enriched cellular components included membrane raft, endosome, apical plasma membrane and focal adhesion. Molecular functions of module 2 were enriched in Protein tyrosine kinase activity, Enzyme binding, Protein binding and Integrin binding (Table 3(b)).

**Table 3:**
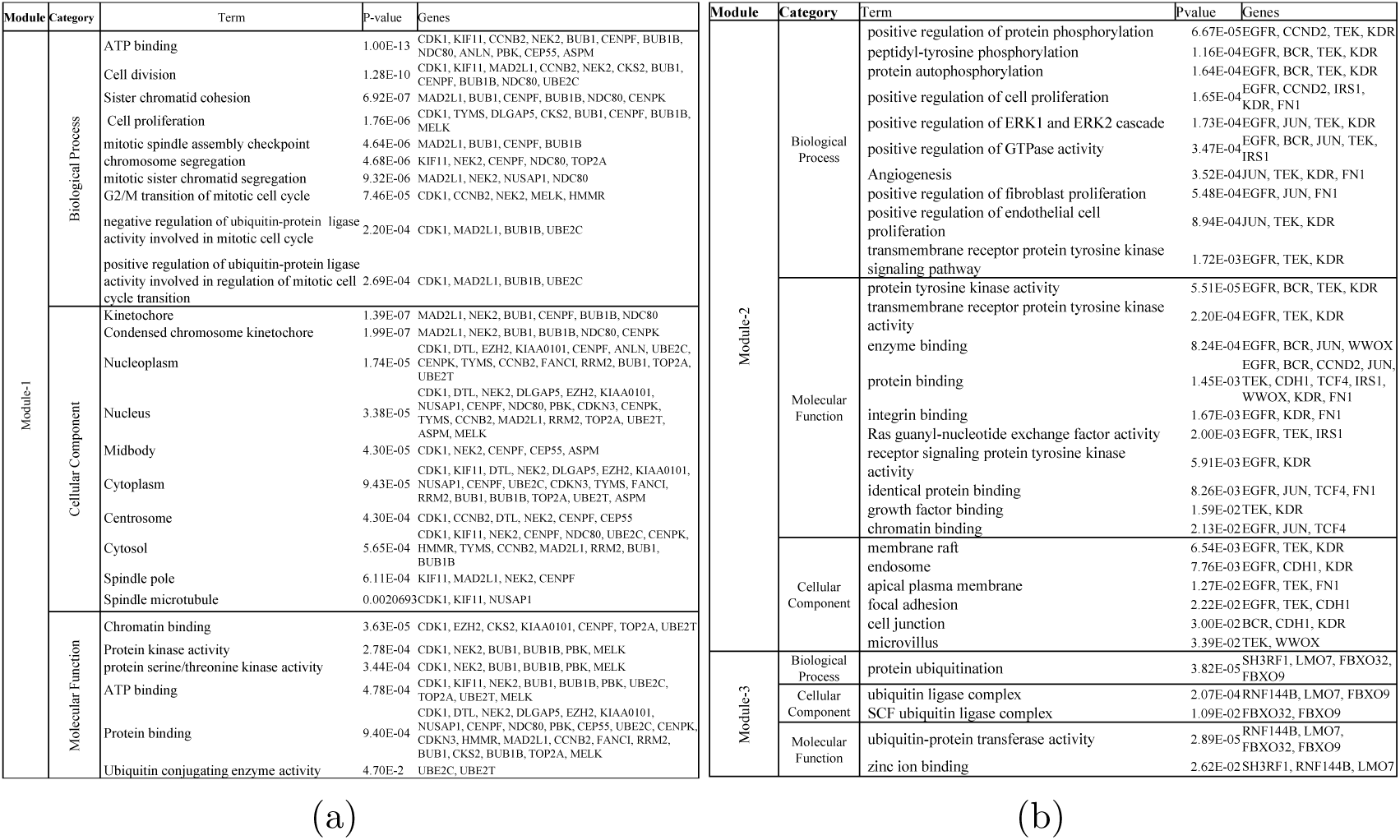
(a) The enrichment analysis of the subnetwork module-1 (b) The enrichment analysis of the subnetworks module-2 and module-3

**Figure 3:**
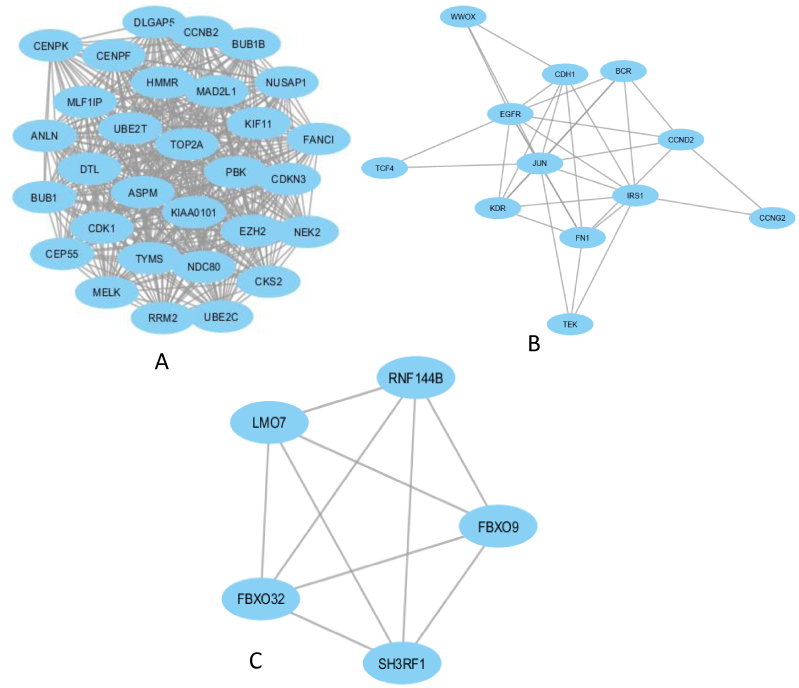
Top three modules in the protein-protein interaction network of the DEGs in TC. The nodes indicate the DEGs and the edges indicate the interactions between two genes.

Module 3 gene ontology annotation found significant enrichment for Protein ubiquitination, with significantly enriched cellular components being ubiquitin ligase complex and SCF ubiquitin ligase complex, which have involvement in proteosomal and autophagy functions. Module 3 molecular functions showed significant enrichment in ubiquitin-protein transferase activity and zinc ion binding (Table 3 (b)). It was also notable that pathway enrichment analyses found modules 1 and 2 were significantly enriched in protein binding pathways, which was not seen in was obtained in module 3 (Table 3 (b)).

### 3.5. Regulatory signatures: TFs and miRNAs affecting DEG transcript Levels

#### 3.5.1. Transcriptional regulatory network construction and TF enrichment analysis

The construction of reporter TFs and DEGs interaction network constructed by NetworkAnalyst revealed a number of potentially important TFs selected by the topological analysis using dual metric approach involving degree and betweenness (Fig. 4). The top 10 ranked TFs with the highest degree and betweenness centrality included FOXC1, GATA2, YY1, FOXL1, E2F1, NFIC, SRF, TFAP2A, HINFP, and CREB1. Pathway enrichment analysis of the TFs that differ between FTC and FTA and DEGs mainly identified as statistically significant a number of relevant pathways that included Transcriptional misregulation in cancer, Acute myeloid leukemia, Pathways in cancer, Thyroid cancer, Cell cycle, Dorso-ventral axis formation, Arrhythmogenic right ventricular cardiomyopathy (ARVC), Cocaine addiction, Prostate cancer pathways (Table 4(b)).

**Table 4:**
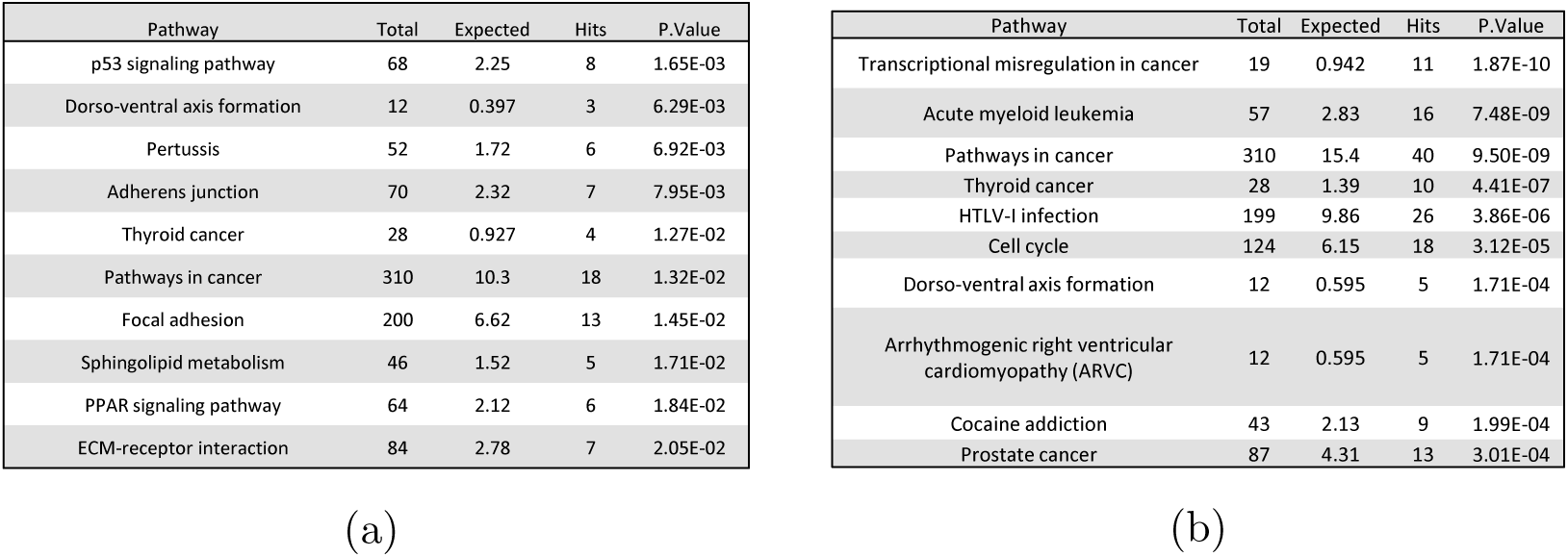
(a) Top 10 KEGG pathway in genes of micro-RNAs and differentially expressed genes interaction networks (b) Top 10 KEGG pathway in genes of TFs and differentially expressed genes interaction networks

| Pathway | Total | Expected | Hits | P.Value |
| --- | --- | --- | --- | --- |
| p53 signaling pathway | 68 | 2.25 | 8 | 1.65E-03 |
| Dorso-ventral axis formation | 12 | 0.397 | 3 | 6.29E-03 |
| Pertussis | 52 | 1.72 | 6 | 6.92E-03 |
| Adherens junction | 70 | 2.32 | 7 | 7.95E-03 |
| Thyroid cancer | 28 | 0.927 | 4 | 1.27E-02 |
| Pathways in cancer | 310 | 10.3 | 18 | 1.32E-02 |
| Focal adhesion | 200 | 6.62 | 13 | 1.45E-02 |
| Sphingolipid metabolism | 46 | 1.52 | 5 | 1.71E-02 |
| PPAR signaling pathway | 64 | 2.12 | 6 | 1.84E-02 |
| ECM-receptor interaction | 84 | 2.78 | 7 | 2.05E-02 |

(a)
| Pathway | Total | Expected | Hits | P.Value |
| --- | --- | --- | --- | --- |
| Transcriptional misregulation in cancer | 19 | 0.942 | 11 | 1.87E-10 |
| Acute myeloid leukemia | 57 | 2.83 | 16 | 7.48E-09 |
| Pathways in cancer | 310 | 15.4 | 40 | 9.50E-09 |
| Thyroid cancer | 28 | 1.39 | 10 | 4.41E-07 |
| HTLV-I infection | 199 | 9.86 | 26 | 3.86E-06 |
| Cell cycle | 124 | 6.15 | 18 | 3.12E-05 |
| Dorso-ventral axis formation | 12 | 0.595 | 5 | 1.71E-04 |
| Arrhythmogenic right ventricular cardiomyopathy (ARVC) | 12 | 0.595 | 5 | 1.71E-04 |
| Cocaine addiction | 43 | 2.13 | 9 | 1.99E-04 |
| Prostate cancer | 87 | 4.31 | 13 | 3.01E-04 |

**Figure 4:**
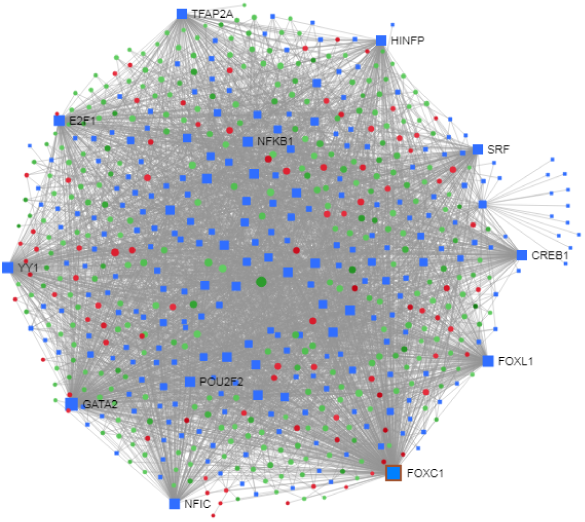
Construction of regulatory networks of TF-DEG interactions. Red nodes indicate up-regulation and green node represents down-regulated DEGs.

#### 3.5.2. miRNA regulatory network construction and enrichment analysis of miRNA

We constructed TC DEG-miRNA interactions networks using Network-Analyst (Fig. 5). By topological analysis, the top 10 ranked miRNAs by highest degree and betweenness centrality were selected which includes hsa-mir-335-5p, hsa-mir-26b-5p, hsa-mir-124-3p, hsa-mir-16-5p, hsa-mir-192-5p, hsa-mir-1-3p, hsa-mir-17-5p, hsa-mir-92a-3p, hsa-mir-215-5p, and hsa-mir-20a-5p. The analysis indicates that these are miRNAs having the highest potential to regulate levels of the DEG transcripts differently in FTC and FTA. The pathway enrichment analysis of the miRNA associated DEGs networks identified the p53 signaling pathway, Dorso-ventral axis formation, Pertussis, Adherens junction, Thyroid cancer, Pathways in cancer, Focal adhesion, Sphingolipid metabolism, PPAR signaling pathway, and ECM-receptor interaction signaling system statistically significant (Table 4(a))

**Figure 5:**
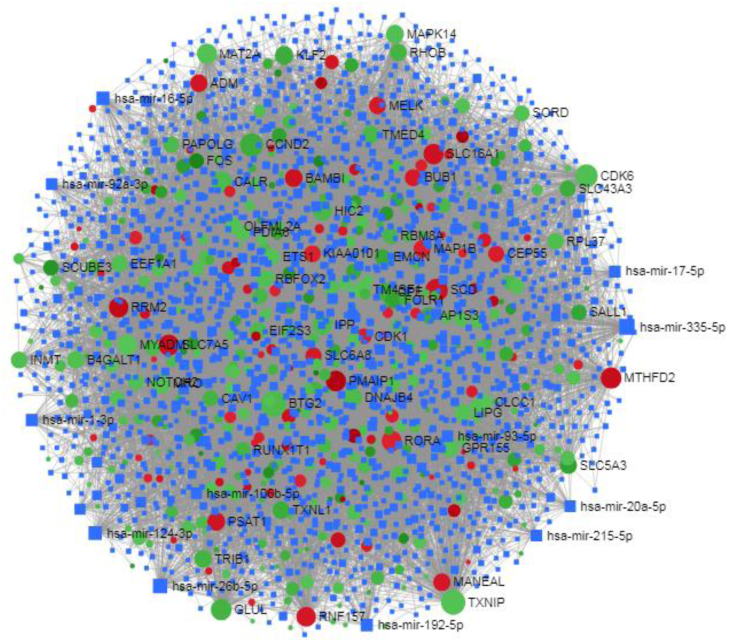
Construction of regulatory networks of DEGs-miRNAs interaction. Red nodes indicate up-regulation and green node represents down-regulated DEGs.

## 4. Discussion

To clarify the molecular mechanisms that underlie the pathological features (especially malignancy potential) that distinguish malignant and benign thyroid lesions we examined differential gene expression between FTC from FTA. Such a comparison is likely to be more informative than making such a comparison of malignant and normal tissues which will contain many more differences related to neoplastic cell development. In addition, the lesions contain significant non-neoplastic cell components (such as mesenchymally-derived and endothelial cells) that are also likely to differ markedly from those of normal tissue.

The DEGs identified from the FTC/FTA comparison were used to identify possible regulatory patterns, key molecular pathways, and protein-protein interactions among these pathway gene products. Such an analytical approach can be used to find molecular signatures that may serve either as potential therapeutic targets or as biomarkers to differentially diagnose or identify FTC. As did the original study by Wojtas *et al.* [51], we identified significantly altered genes that may be candidate biomarkers that distinguish FTC. However, another important use of such data (and our main focus here) is to identify and characterize biological functions associated with these genes that may give insights into the biology and behaviour of FTC lesions themselves.

Our study thus identified a number of important pathways that were differentially expressed in FTC, some of which would be expected by the malignant profile of FTC compared to FTA. For example, the p53 signaling pathway plays an important role in cancer cell apoptosis and DNA repair by causing cycle arrest in response to DNA damage [57] and previously implicated in thyroid cancers [12]. Cell cycle progression is regulated and facilitated by cyclin-dependent kinases that are activated by cyclins such as cyclin D1. Indeed, cyclin D1 levels can influence tumour progression, and may have prognostic significance in FTC [50]. Related to this, it has also been reported that the One-carbon metabolism pathway is actively involved in cancer progression, probably due to its involvement in nucleotide synthesis [33].

We also found a number of significant GO groups are associated with the DEGs as shown in table 2. PPI network reconstruction and the analysis of reconstructed PPI network represent a powerful approach for understanding disease mechanisms, so to construct a PPI network for the DEGs in our study we combined results of statistical analyses with the protein interactome network. To identify potential hub proteins, topological analysis strategies were employed. This identified 12 hub genes (Table 1(b)). These include a number of genes commonly associated with cancers (including thyroid cancers), such as EGRF, JUN, FOS, CDK1 and other cyclin pathway genes, E-cadherin (CDH1) and E2F1. The hub protein EGFR is linked to growth in many types of cancer as well as cardiovascular diseases, and has been previously linked to thyroid cancers [42]. JUN and FOS are best characterised as subunits of the AP-1 transcript factor that has a crucial role in many types of cell differentiation and inflammation processes [6]. GO databases also delineated particular roles for JUN in Angiogenesis and regulation of Endothelial cell proliferation, both of which have central roles in metastasis and invasion (Table-2 (b)). CDK1 is an important cell cycle regulator and is involved in the breast, lung and ovary carcinomas [26]. CDK1 overexpression has been documented in lung cancer, lymphoma and advanced melanoma while loss of cytoplasmic CDK1 predicts poor patient survival and may confer chemotherapeutic resistance in the latter [36]. Cyclin B1 (in the same family as CDK1) overexpression and/or mislocalization has been described in several primary cancers including thyroid carcinoma, colon, gastric, prostate, breast, and non small-cell lung cancer [36]. It is also actively involved in mitotic cell cycle phase transition (Table 2 (a)). CDKN3 encodes a cyclin inhibitor (regulating cell cycle) and has been described as overexpressed in lung adenocarcinoma (ADC), squamous cell carcinoma (SCC), hepatocellular carcinoma, cervical cancer and epithelial ovarian cancer [56]. Over-expression of EZH2 is frequently observed in many cancer types [54].

TYMS expression is also associated with the risk of development of epithelial cancers and lymphoma cancer[19]. Besides, in Hereditary diffuse gastric cancer (HDGC), the hub gene CDH1 mutations are connected with an increased incidence of lobular carcinoma of the breast and, possibly, prostate cancer and colorectal carcinoma [35]. This gene is also involved in extra-cellular matrix organization (Table 2(b)). Hub gene CCNB2 is known to be underexpressed in thyroid cell tumors [10]. We found that CDK1 and CCNB2 hubs genes were actively associated with p53 signaling pathway, Progesterone-mediated oocyte maturation and Cell cycle pathways. The hub gene TYMS is also involved in One carbon pool by folate pathway, while FOS, JUN and CDH1 genes are involved in Pathways in cancer and JUN, FOS genes in Estrogen signaling pathway. A number of other hub genes have been described as associated with malignant thyroid cancers including EXH2, UBCH10,TFAP2A, TOP2A and SRF [27, 23, 8, 41, 21]. By considering the possible roles of these hub proteins in pathogenesis of FTC and other related diseases new roles for these proteins may be identified. In addition to the above hub genes, we also identified a number of factors not previously noted as having a role in FTC or thyroid cancers, including UBA52, GATA2, FOXL1 and NFIC.

As we know that the regulation of gene expression is controlled by TFs and miRNAs at post-transcriptional and/or transcriptional levels, so changes in these molecules may provide crucial information regarding dysregulation of gene expression in FTC. Thus, we investigated TFs and their relationship to DEGs in these tumours. One of the TFs, we identified was FOXC1 which a previous study found that FOXC1 was strongly associated with thyroid cancers [5]. Overexpression of YY1 in differentiated thyroid cancers has also been noted [2]. The pathway analysis of these TFs showed statistical significance in FTC. From the DEGs a miRNA interaction network of miRNAs (Table 1(b)) was also identified and analysed. Our pathway analysis of these miRNA also showed statistically significant relationships with FTC. Notably, the target DEGs of these miRNAs and TFs included the p53 signaling pathway, PPAR signaling pathway, Dorso-ventral axis formation, Focal adhesion (Table 4(a)) and Pathways in cancer, Transcriptional misregulation in cancer, and Thyroid cancer, respectively. The miRNA species we identified that have previously been shown to have a role in thyroid cancer included hsa-mir-335-5p, hsa-124-3p, hsa-mir-17-5p and hsa-mir-20a-5p [9, 55, 25, 17]. However, 6 other miRNAs we identified have not previously been associated with thyroid lesions. miRNA species have wide ranging effects on gene expression which may be indirect, so further analysis for these are needed.

We analysed the dataset originally generated by Wojtas *et al.*, who sought markers distinguishing FTC and FTA and combined microarray methods and literature meta-analysis to achieve this. The gene markers they identified included TFF3 and CPQ (previously characterised FTC markers), PLVAP and ACVRL1, all genes expressed in thyroid although ACVR1 is an activin receptor with wide expression. These latter genes were not identified by our analytical approach, but it should be noted that our study used very different analytic tools, focussing on identifying pathways and on finding evidence for gene functions using PPI and other resources. Our analytical approach was somewhat more similar to that of Wang *et al.* [49] who examined an older thyroid cancer dataset (GSE27155) containing 10 FTA, 14 FTC, 4 normal thyroid tissue samples plus other types of thyroid lesion. While we employed a number of additional analytical tools (some developed previously by us) there were a number of significant genes and pathways we found in common in the FTC/FTA analyses Wang *et al.* [49]. These notably include the AP-1 TF and its components (e.g., JUN and FOS), BCL2, factors relating to DNA damage and cyclin-related factors that regulate cell cycle. We identified no genes that Wang *et al.* identified in comparing FTC and normal tissues. Recent studies by Shang *et al.* and by Liang and Sun [24] examined datasets generated from papillary thyroid cancer lesions that were compared with and normal tissues. DEGs identified in these studies were analysed by methods related to our approach [44]. However, unlike the Wang *et al.* study cited above, very few of the hub genes these studies identified were found in our analysis; BCL2 was the main exception. This lack of concordance with the FTC studies may well reflect the different tumour type (and lower malignancy rates) represented by papillary thyroid cancer, and it might be useful to compare all these datasets in detail to determine how these types of tumour differ.

## 5. CONCLUSIONS

Our data-driven approach has uncovered a number of significant molecular mechanisms that may underlie the difference between FTC and FTA. In this study, using integrated bioinformatics analyses, we analyzed gene expression profiles in FTC and FTA and used this information to identify candidate hub proteins that could play significant roles in FTC development. Our results indicate the directions for future experimental work needed to clarify the roles of these cellular proteins. In sum, the study identified gene networks that advance our understanding of the pathogenesis of FTC, and indicates new avenues to develop therapies for FTC.

## Acknowledgments

The work was supported by the iHealthOmics (http://www.ihealthomics.com/) and a University of Sydney Fellowship award.

## Author Contribution

conceptualization, MAH and MAM and TAA; methodology, MAH and TAA.; software, MAH; validation, MAH, TAA and MAM; formal analysis, MAH; investigation, MAH; resources, MAH; data curation, MAH and TAA; writingoriginal draft preparation, MAH; writingreview and editing, MAH, MMR, JQ and MAM; visualization, MAH and TAA; supervision, MAM; project administration, FH and MAM; funding acquisition, FH and MAM

This research received no external funding

We have No conflict of interest.

### The following abbreviations are used in this manuscript

ADC: Adenocarcinoma
DEG: Differencially Expressed Genes
FC: fold change
FTC: Follicular Thyroid Carcinomas
FTA: Follicular Thyroid Adenoma
GO: Gene Ontology
GEO: Gene Expression Omnibus
HDGC: Hereditary diffuse gastric cancer
KEGG: Kyoto encyclopedia of genes and genomes
miRNA: Micro-RNA
PPI: Protein-protein Interactions
SCC: Squamous Cell Carcinoma
TF: Transcription Factor

